# *W*_*d*_^*^-test: Robust Distance-Based Multivariate Analysis of Variance

**DOI:** 10.1101/492413

**Authors:** Bashir Hamidi, Kristin Wallace, Chenthamarakshan Vasu, Alexander V. Alekseyenko

## Abstract

**Background:** Community-wide analyses provide an essential means for evaluation of the effect of interventions or design variables on the composition of the microbiome. Applications of these analyses are omnipresent in microbiome literature, yet some of their statistical properties have not been tested for robustness towards common features of microbiome data. Recently, it has been reported that PERMANOVA can yield wrong results in the presence of heteroscedasticity and unbalanced sample sizes.

**Findings:** We develop a method for multivariate analysis of variance, 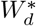, based on Welch MANOVA that is robust to heteroscedasticity in the data. We do so by extending a previously reported method that does the same for two-level independent factor variables. Our approach can accommodate multi-level factors, stratification, and multiple *post hoc* testing scenarios. An R language implementation of the method is available at https://github.com/alekseyenko/WdStar.

**Conclusion:** Our method resolves potential for confounding of location and dispersion effects in multivariate analyses by explicitly accounting for the differences in multivariate dispersion in the data tested. The methods based on 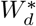 have general applicability in microbiome and other ‘omics data analyses.

## 1 Introduction

Beta diversity analyses or community-wide ecological analyses are important tools for understanding the differentiation of the entire microbiome between experimental conditions, environments, and treatments. For these analyses, specialized distance metrics are used to capture the multivariate relationships between each pair of samples in the dataset. Analysis of variance-like techniques, such as PERMANOVA [1], may then be used to determine if an overall difference exists between conditions. The distances use all of the measured taxa information simultaneously without the need to explicitly estimate individual covariances. The utility of these methods is hard to underestimate as virtually every recent major microbiome report has used some form of a community-wide association analysis. On many occasions the comparison reveals major differences between the groups. However, one is not guaranteed to find one. For example, in Redel et al. [2] the authors have found that there are significant differences in cutaneous microbiota in diabetic vs. non-diabetic subject feet, but not on their hands (see figure 5). This lack of difference is an important indicator about the potential pathobiological processes that lead to diabetic foot ulcers. Therefore, getting the correct result in such comparisons is important.

From the statistical stand point, community-wide analyses test the hypothesis that the data from two or more conditions share the location parameter (centroid or multivariate mean). Caution, however, needs to be taken to ensure that potential violations of assumptions do not lead to adverse statistical behavior of PERMANOVA. Two such assumptions that are commonly violated are the multivariate uniformity of variability (homoscedasticity) and sample size balance. We have previously shown that simultaneous violation of both assumptions leads to PERMANOVA analysis with indiscriminate rejection and type I error inflation or to significant loss of power up to inability to make any rejections at all [3]. Unfortunately, heteroscedasticity across conditions is a very common feature of microbiome data. Thus new robust methods are needed to ensure correct data analysis.

We have previously described a 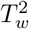 test, which presents a robust solution for comparing two groups of microbiome samples [3]. The two-sample scenario is common, but not universally satisfying as many study designs often include many different sample types, e.g. from affected and unaffected sites of a study subject and from a matched healthy control [4]. Here we describe a further extension of 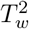 to allow for arbitrary number of groups with possibly different within group variability to be compared using an omnibus test for equality of means. Our method presents an advance to the state-of-the-art by introducing a way to compare data from multiple conditions where heteroscedasticity is a nuisance and only the differences between location of the data are important.

## 2 Univariate Welch MANOVA

Univariate solutions for a heteroscedastic test to compare *k*-means deal with finding asymptotic distributions for 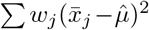, as defined later in equations (2) and (3). Welch’s solution [5] is perhaps the most known and well adopted in statistical literature. Next we briefly describe it, as we will build on extending this statistic to multivariate data.

Suppose we observe data from *k* populations 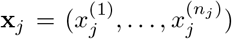 with potentially unequal number of observations, *n_j_* for *j* = 1, …, *k*, in each. Let 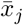 and 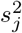 denote the means and variances for each sample. The Welch ANOVA statistic is

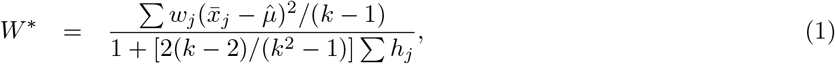

where

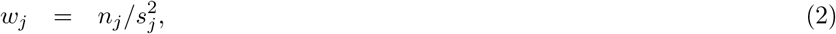

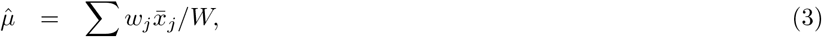

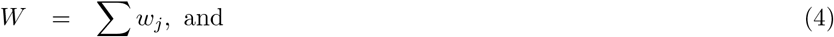

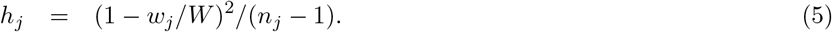

The Welch test uses *F*(*k* – 1, *f*), for *f* = (*k*^2^ – 1)/(3/ ∑ *h_j_*) distribution to draw inference with *W*\*[5].

## 3 Calculation of multivariate Welch W-statistic on distances

To derive a Welch *W*\* statistic suitable for analysis of microbiome data, 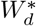 we follow the same approach as we did in our derivation of 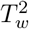. We first demonstrate that in the univariate case 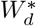 can be expressed in terms of sums of pairwise square differences. Next we observe that these sums represent the squares of the univariate Euclidean distances, which allows for a direct extension of the 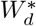 statistic computation for multivariate Euclidean distances and in fact any arbitrary distance or dissimilarity metric. The derivation of the statistic in terms of dissimilarities makes it suitable for analysis of microbiome data via a permutation test.

We have previously shown [3] that the sample variances can be written as

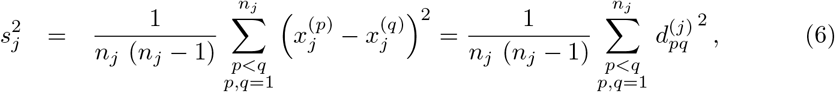

where 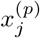 and 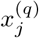 denote *p*-th and *q*-th observations in the *j*-th level, 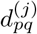 is distance between them. Hence,

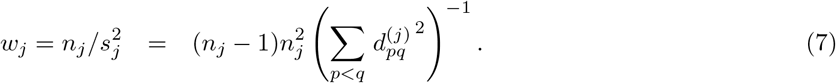

Now consider,

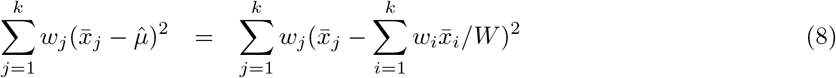

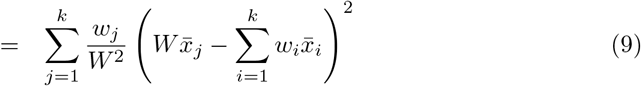

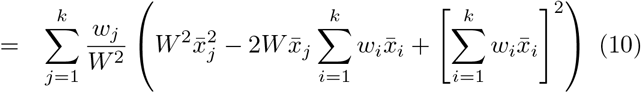

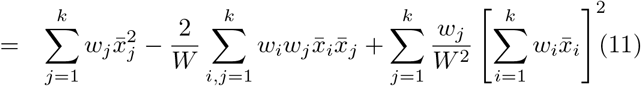

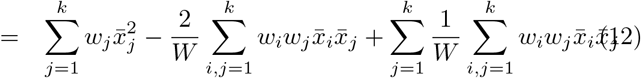

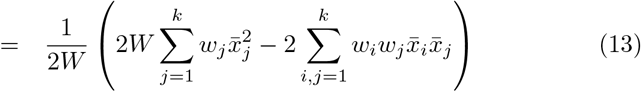

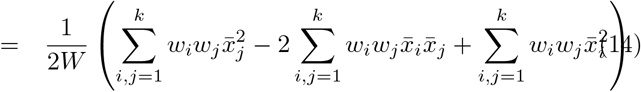

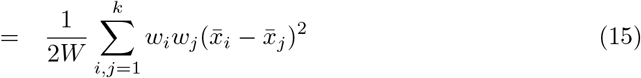

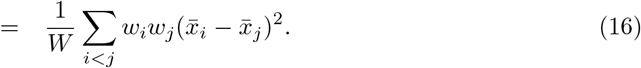

Equation (16) means that 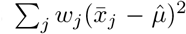 can be expressed as weighted sum of squares of pairwise inter-group mean differences, which makes for a convenient expression to compute. Finally, we have previously shown that squares of mean differences can be expressed in terms of squares of pairwise sample differences [3], i.e.

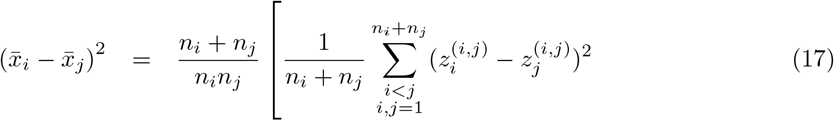

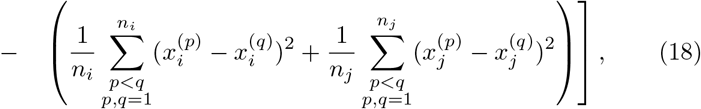

where 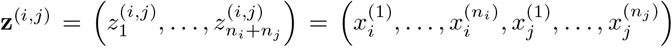. The squares of the pairwise differences under the summations in equation (18) can be thought of as the squares of the pairwise Euclidean distances in one dimension. This allows us to generalize the univariate Euclidean Welch ANOVA to MANOVA with arbitrary distances, where the distances can be suitably defined for the data at hand, including all of common distances used with microbiome data.

Note that in contrast to the PERMANOVA statistic, the distance-based 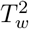 and 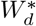 explicitly account for potentially unbalanced number of observations and differences in multivariate spread in the two samples. Finally, observe that 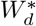 reduces to 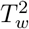 when *k* = 2, as *W*\* reduces to Welch t-statistic.

As with 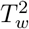, the exact distribution of the multivariate distance-based 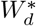 statistic is dependent on many factors, such as the dimensionality of underlying data, distributions of the random variables comprising the data, the exact distance metric used, and the number of groups compared *k*. To make a practical general test, we use permutation testing to establish the significance. To do so, we compute 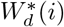 on *m* permutations of the original data, for *i* = 1, …, *m*, and estimate the significance as the fraction of times the permuted statistic is greater than or equal to *W_d_*, i.e. 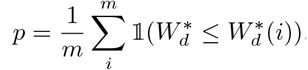. Here 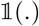 designates the indicator function.

Confounder modeling and repeated measures are often key elements of microbiome study design. These can be accounted for in permutation testing procedures using restricted permutation. For example, the effect of a discrete valued confounder can be removed from the P-value calculation by restricting permutations to only within the levels of the confounding variable. This amounts to an application of stratified analysis of variance. Similarly, restricting permutations to within individual subjects only, results in a repeated measures analysis. Notice that the test statistic under restricted permutations remains the same, but the null distribution is changed to reflect the desired comparison. Methods for 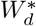 and these restricted permutation methods are implemented in our software, available at https://github.com/alekseyenko/WdStar.

When multiple means are compared with 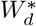 a statistically significant result may prompt the question about attribution of the differences to a specific group or groups. *Post hoc* testing procedures are used to perform that kind of analysis. There are many possible ways to design the *post hoc* testing procedures, but the guiding principle due to potential for loss of power to multiple testing should be to minimize the number of tests performed. For this reason, in addition to all possible pairwise (one versus one) tests, it may be interesting and relevant to test one group versus all others. In this scenario, samples from one experimental group are compared to pooled samples from the remaining groups. The statistical test for this comparison can equivalently be either 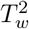 or 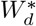 on two level factors. We illustrate the use of one versus all *post hoc* procedure in our application example in section 5 and provide corresponding computation routines in our software.

## 4 Empirical evaluation of 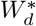 type I error

The principal evaluation that is required to assure statistical properties of 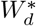 is demonstration of appropriate type I error control. For this purpose, we consider the univariate heteroscedastic case with 3 groups, 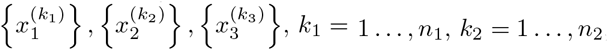, and *k*_3_ = 1 …, *n*_3_, of samples to compare, where *n*_1_, *n*_2_, *n*_3_ are the numbers of observations in each group. We let 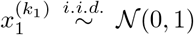 be the reference group, and 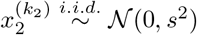 and 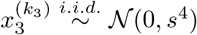 be the groups with different variance *s*^2^ and *s*^4^, respectively, to introduce heteroscedasticity. In our simulation, we let *s*^2^ = {1,0.8,0.2} to control the degree of heteroscedasticity in the range from none to large. Finally, we let the sample sizes *n*_1_, *n*_2_, and *n*_3_ take values of 5, 10, 20, or 40 to generate data with varying total sample size and degree of balance. For each combination of sample sizes and variance we have performed 1,000 simulations of the data for a total of 192,000 datasets. Each dataset has been analyzed using our reference implementation of 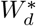, PERMANOVA (adonis function in R library vegan), and univariate Welch ANOVA (oneway.test in R library stats). For distance-based methods, Euclidean distances have been used. Details of simulation are available as a knitted R Markdown file in Additional File 1.

The simulation results comprise the fraction of rejected null hypotheses at *α* = 0. 05 by each test (Figure 1A). A test properly controlling the type I error is expected to have the fraction of rejections equal to the nominal error rate (0.05). Notice that the proposed 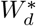 test, in fact, produces the expected error rates over the entire range of simulation parameters. Similarly to our previous observations in the two-sample case, PERMANOVA is not robust to heteroscedasticity when sample size imbalance is present. Observe that whenever the number of observations in the reference group (the one with variance equal to 1) is smaller than that in the less dispersed groups the fraction of rejections is overly inflated, resulting in higher type I error. Also notice that when there are more observations in the reference group than in others (e.g. *n*_1_ = 40, *n*_2_, *n*_3_ < 40) it is hard for PERMANOVA to make the rejections, resulting in approximately zero type I error.

**Figure 1.**
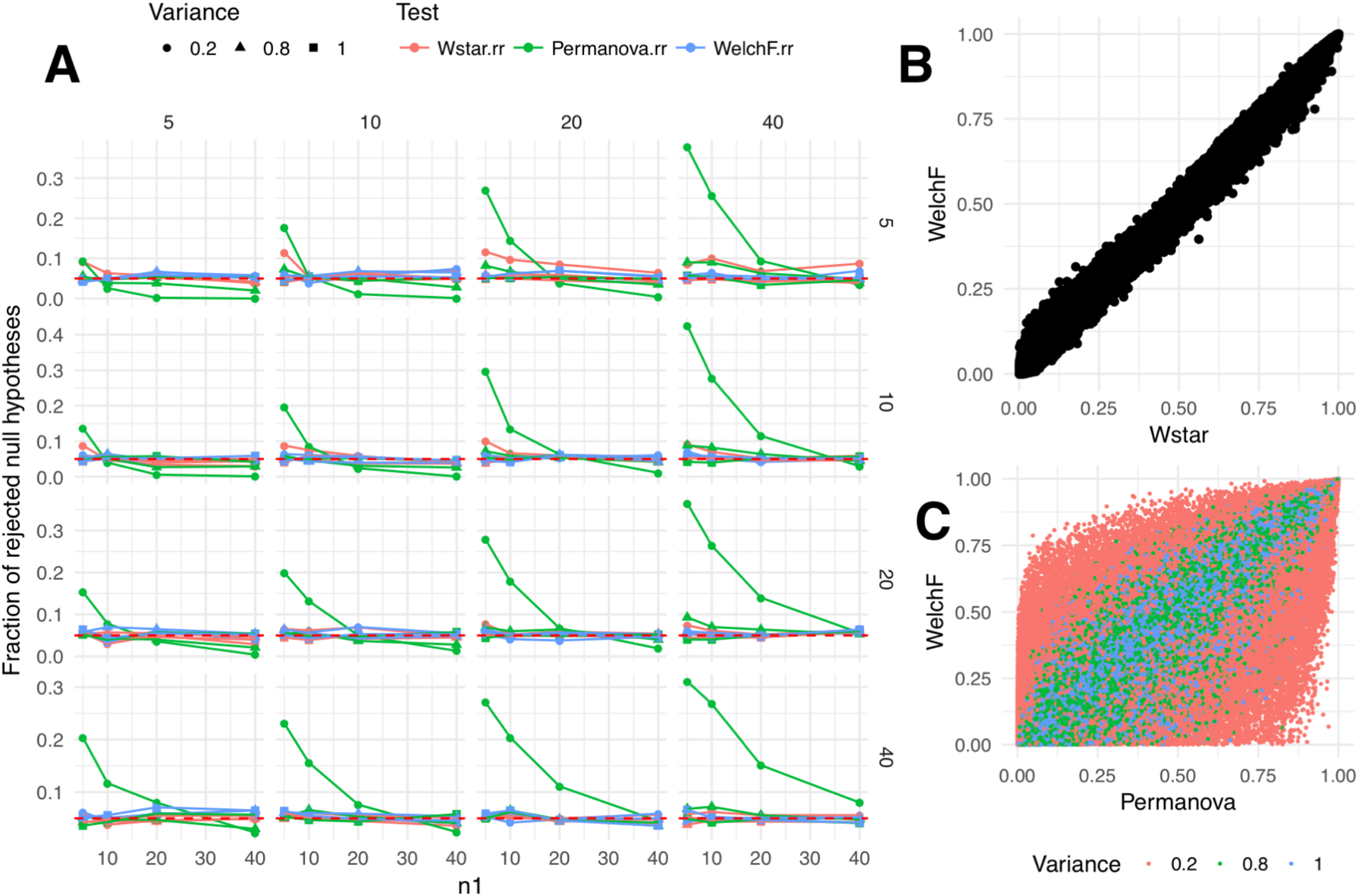
Evaluation of type I errors of 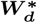 and PERMANOVA permutation tests. Simulation under the null hypothesis results for comparison of 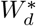 (Wstar), PERMANOVA (Permanova) and distribution-based Welch ANOVA F (WelchF) tests are presented. In panel A, we evaluate the fraction of null hypotheses that have been rejected by each test at *α* = 0.05. The subpanels of A, correspond to simulated datasets with corresponding number of samples in the non-reference groups, with columns corresponding to the least dispersed and rows corresponding to the most dispersed sample. In panel B, the raw p-values from 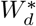 test are plotted against those for the same data with Welch ANOVA F-test. In panel C, we do the same for PERMANOVA p-values and color the points by respective degree of heteroscedasticity in the simulated dataset.

Interestingly, when we compare the raw p-values obtained from 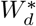 to those from the distribution based asymptotic Welch test, we see a good concordance between the two (Figure 1B). The variability around the trendline is most likely due to Monte Carlo error associated with permutation testing and small sample size. On the contrary, when PERMANOVA is compared to the distribution-based asymptotic test the fit is clearly much noisier (Figure 1C). The concordance is much smaller for tests involving groups with larger degree of heteroscedasticity. The code used to produce the plots in Figure 1 is available as Additional File 2.

Finally, given the equivalence of the 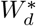 to 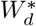 for *k* = 2, and the fact that the two-level test is powered similarly to PERMANOVA, we expect the test described in this paper to be of similar power for *k* > 2 as well. The full empirical evaluation of power characteristics for *k* > 2 is hard to achieve in non-superficial setups as most realistic simulation scenarios present an infinite universe for choice of parameters.

## 5 Application example: Colorectal cancer disparity and microbiome

Extensive scientific literature suggests and important, yet not fully understood role of the intestinal microbiome in the development, progression, and treatment of colorectal cancers (CRC). Several genus level bacterial taxa have been associated with CRC [6] but the role of personal characteristics in influencing the presence of CRC-associated bacteria is not well understood. A few studies have noted marked differences in the microbial environment in the gut of AAs versus others [6, 7, 8, 9, 10] and suggested differences in microbial composition among those with and without colorectal polyps and cancer. Others found distinct differences in the microbes populating the proximal and distal colo-rectum [11, 12]. Lower socioeconomic status and western diet have been associated with a lower microbial diversity, especially in the distal colon [13, 14]. Microbial signature approaches have been used for development of diagnostic biomarkers [8, 15, 16, 17] or assessing differences in immune gene expression [12] – highlighting the increasing importance of statistical methods to analyze clusters of microbes-genes while also taking into account patient level variables. The role of the gut microbiome in CRC disparities is likewise poorly understood [18]. Here we use a pilot CRC dataset to demonstrate the utility of 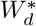 in uncovering signals potentially missed due to heteroscedasticity.

The Medical University of South Carolina (MUSC) Institutional Review Board approved all study activities. The Cancer Registry at Hollings Cancer Center (HCC) at MUSC was used to identify all cases of CRC. The study population was comprised of a sample of histologically-confirmed cases diagnosed between January 1, 2000 and June 30, 2015. Patients were of either AA or CA descent. We abstracted data on demographic characteristics, clinical and pathological variables at diagnosis, treatment received, and patient outcome from the cancer registry. For each case, we also obtained a formalin-fixed, paraffin-embedded tissue blocks from the MUSC Department of Pathology and Laboratory Medicine. DNA was extracted following standard protocols in the laboratory. Briefly, the colonic tissue was transferred to a tube containing lysis buffer (1% SDS, 1 mg/ml Proteinase K, LTE pH 8.0). The solution was incubated at 50°C for 1 hour, followed by phenol/chloroform extraction and ethanol precipitation. The quantity and quality of DNA was then determined by running a small aliquot on a 1% agarose gel and comparing it to a set of DNA standards. The extracted DNA was stored at -80°C. V3 and V4 regions of the 16S rRNA gene have been amplified using 16S Amplicon PCR Forward Primer = 5’ TCGTCGGCAGCGTCAGATGTGTATAAGAGACAGCCTACGGGNGGCWGCAG 16S Amplicon PCR Reverse Primer = 5’ GTCTCGTGGGCTCGGAGATGTGTATAAGAGACAGGACTACHVGGGTATCTAATCC using KAPA HiFi enzyme. The library has been prepared using Nextera XT index kits, and sequenced using MiSeq Reagent Kit v3 in a Miseq instrument. Taxonomic assignments have been generated using QIIME preprocessing application of Illumina Basespace platform with default parameters. Using genus level data restricted to genera previously reported in a systematic review to be associated with CRC [19], Jensen-Shannon Divergence distances have been computed between the subjects of Caucasian and African American races with cancers in distal and proximal locations of their colons (Table 1). See Additional file 4 for the list of 14 genera retained for this analysis.

**Table 1.** Number of the subjects in the colorectal cancer example analysis.

| Race | Cancer Location | N |
| --- | --- | --- |
| African American | distal | 2 |
|  | proximal | 3 |
| Caucasian | distal | 5 |
|  | proximal | 4 |

We selected a convenience sample from our MUSC cancer cohort of 20 patients (10 AAs, 10 CAs) which we matched on colonic location (proximal, distal) and sex. Of the 20 cases, 6 have been removed due to low sequence count (< 100) within the genera of interest. Due to extremely small pilot-scale sample size, the group unbalance and potential for heteroscedasticity prompt caution with using PERMANOVA for these comparisons (Figure 2). Indeed, the race and location interaction model achieves significance (*P* < 0.05) with 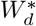 test, while the PERMANOVA result is insignificant (*P* = 0.28) (Table 2). Likewise, there is a discrepancy in test results for the primary effect of the race at 0.05 significance threshold.

**Figure 2.**
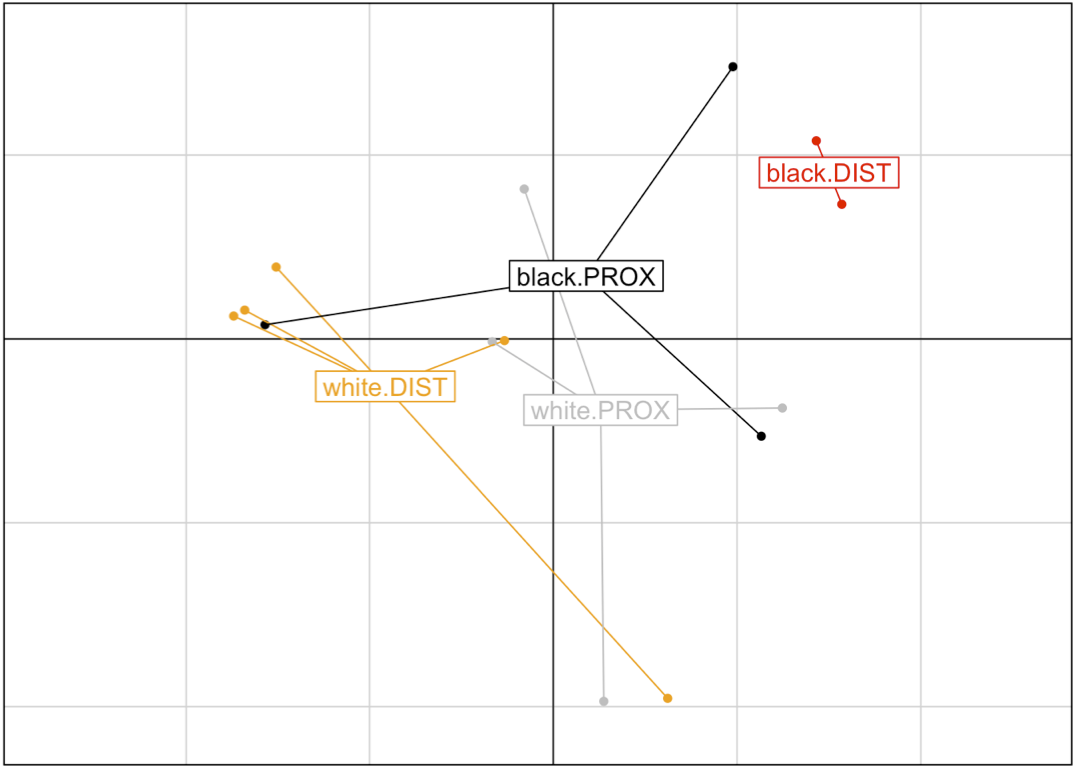
PCoA plot of the JSD distances between CRC microbiome samples. African American distal (red) samples appear to be separated on PC1 from the samples in the proximal AA (black) and Caucasian (gray) and Caucasian distal (orange) samples. Likewise, the plot suggest that the multivariate spread may differ dramatically in the compared groups with AA distal samples being most concentrated relative to the other groups.

**Table 2.** Significance of the primary and interaction effects by PERMANOVA and 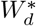 tests.

| Covariate | PERMANOVA P-value | $W_d^*$ P-value |
| --- | --- | --- |
| Race | 0.064 | 0.047 |
| Location | 0.907 | 0.908 |
| Race & Location | 0.282 | 0.037 |

Significance of the interaction term may dictate additional questions about, which groups differ from the rest. We demonstrate the use of one versus all *post hoc* testing by comparing each group with the rest of the samples (Table 3). As expected, these indicate a significant difference (*P* < 0.05) in the microbiome of the African American distal CRC samples from the rest, and a trend for difference of the Caucasian distal samples. Note that the interpretations of these results might differ if multiple comparison issues are taken into account. Due to the pilot nature of these data, we do not perform any formal corrections, as our goal is to determine the plausibility of significant differences, which are to be evaluated in appropriately sized datasets where power is not a concern.

**Table 3.** One versus all *post hoc* comparisons of the interaction terms.

| Group | $T_w^2$ statistic | $W_d^*$ P-value |
| --- | --- | --- |
| AA distal | 8.88 | 0.039 |
| CA distal | 1.93 | 0.075 |
| AA proximal | 0.36 | 0.936 |
| CA proximal | 0.70 | 0.665 |

The data and R Markdown for this application is included in Additional file 4.

## 6 Discussion and Conclusion

Community-wide analyses where the entire microbiome is modelled as a response variable of one or more factors has become a standard first-line of analysis technique in the field. These techniques address the question of overall aggregate changes in the microbiome in response to explanatory variables without the need to model each individual microbiome constituent. PERMANOVA [1] has been one of the most dominant tools for such analyses, although the potential for confounding of location and dispersion effects has been recognized for a long time [20, 21]. The 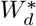 method closes the gap by explicitly accounting for the differences in multivariate dispersion in the data tested, which has been shown to be associated with adverse statistical properties in PERMANOVA [3]. Current heteroscedasticity-aware methodologies allow for modeling multi-level factors, stratification, and multiple *post hoc* testing scenarios.

Although originally developed for discrete-valued covariates, PERMANOVA remains a viable analysis option for continuous covariates as well when multivariate regression-like formula are utilized [22]. However, the effect of heteroscedasticity has not been rigorously evaluated or addressed for such analyses. To be fair, heteroscedasticity with continuous covariates is an issue that does not have a generic statistical solution applicable in most cases. A more cautious analysis involving continuous covariates may require corroboration with discretized independent variables by 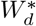, but has to also account for potential statistical power issues pertaining to discretization.

A major limitation of most community-wide analyses is that those often do not yield a natural unified framework for evaluation of taxon-level effects. Currently, methods that have this unifying ability are emerging [23]. None of these, however, are evaluated for robustness with heteroscedastic data yet.

## Supporting information

## Ethics approval and consent to participate

The human subjects component of this research has been approved by the Medical University of South Carolina (MUSC) Institutional Review Board.

## Consent for publication

Not applicable.

## Author’s contributions

AVA has conceived the method, derived the test statistic, and developed reference implementation in R statistical programming language, wrote the manuscript and performed data analysis; BH has implemented code for restricted permutations; KW has designed original study on CRC, and collected and organized tissue and DNA samples; CV has generated 16S rRNA gene sequencing data. All authors have reviewed and approved the manuscript.

## Acknowledgements

The authors would like to thank ZhengZheng Tang for early input in this work.

## Additional Files

Additional file 1 — Test_Wstar_simulation.html

Knitted HTML R Markdown document detailing the steps of producing the simulation datasets and running each test to evaluate the Type I error performance of 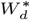 relative to PERMANOVA and asymptotic Welch F test.

Additional file 2 — plot_Wstar.html

Knitted HTML R Markdown document containing the code used to produce Figure 1.

Additional file 3 — MUSC_CRC.RData

R Data file containing the R package phyloseq object with data for the application example. The object includes the genus level abundance tables, sample data containing designations of the race and CRC location, and taxonomic table for the data.

Additional file 4 — 16S_alone_taxa_of_interest.html

Knitted HTML R Markdown document detailing application example analyses.

## Availability of data and materials

All data, software and other materials are available at https://github.com/alekseyenko/WdStar.

