## Supplementary material for "*W*_*d*_^*^-test: Robust Distance-Based Multivariate Analysis of Variance"

CRC µbiome


### CRC µbiome

###### *Alexander V. Alekseyenko*

###### *Thu May 31 09:03:29 2018*

### Load the data

```
library(phyloseq); packageVersion("phyloseq")
```

```
## [1] '1.23.1'
```

```
load("MUSC_CRC.RData")
physeq
```

```
## phyloseq-class experiment-level object
## otu_table()   OTU Table:         [ 306 taxa and 14 samples ]
## sample_data() Sample Data:       [ 14 samples by 2 sample variables ]
## tax_table()   Taxonomy Table:    [ 306 taxa by 7 taxonomic ranks ]
```

```
unique(tax_table(physeq)[,"Genus"])
```

```
## Taxonomy Table:     [14 taxa by 1 taxonomic ranks]:
##         Genus             
## 4372795 "Fusobacterium"   
## 981783  "Staphylococcus"  
## 261924  "Ruminococcus"    
## 592925  "Prevotella"      
## 2949328 "Bacteroides"     
## 274016  "Lactobacillus"   
## 179814  "Akkermansia"     
## 183157  "Faecalibacterium"
## 71638   "Porphyromonas"   
## 813479  "Bifidobacterium" 
## 345542  "Roseburia"       
## 606755  "Dysgonomonas"    
## 835900  "Odoribacter"     
## 565720  "Treponema"
```

### Visualization

#### compute distances

```
# Jensen Shannon distances for microbiome data
phy.dist = phyloseq::distance(physeq, method="jsd")
```

```
library(ade4)
library(vegan)
```

```
## Loading required package: permute
```

```
## Loading required package: lattice
```

```
## This is vegan 2.5-2
```

```
phy.pco = dudi.pco(phy.dist, scannf=F, nf=3)
```

```
## Warning in dudi.pco(phy.dist, scannf = F, nf = 3): Non euclidean distance
```

```
race.location = with(sample_data(physeq), interaction(race, location))
meandist(phy.dist, race.location)
```

```
##            black.DIST white.DIST black.PROX white.PROX
## black.DIST  0.1181524  0.6605857  0.4460758  0.5677946
## white.DIST  0.6605857  0.5469615  0.6033687  0.6035597
## black.PROX  0.4460758  0.6033687  0.6880283  0.6175192
## white.PROX  0.5677946  0.6035597  0.6175192  0.6596153
## attr(,"class")
## [1] "meandist" "matrix"  
## attr(,"n")
## grouping
## black.DIST white.DIST black.PROX white.PROX 
##          2          5          3          4
```

```
diag(meandist(phy.dist, race.location))
```

```
## black.DIST white.DIST black.PROX white.PROX 
##  0.1181524  0.5469615  0.6880283  0.6596153
```

```
s.class(phy.pco$li, race.location, cellipse = 0, 
        col = c('red', 'orange', 'black', 'gray75'))
```

```
adonis(phy.dist~race.location, permutations=99999)
```

```
## 
## Call:
## adonis(formula = phy.dist ~ race.location, permutations = 99999) 
## 
## Permutation: free
## Number of permutations: 99999
## 
## Terms added sequentially (first to last)
## 
##               Df SumsOfSqs MeanSqs F.Model      R2 Pr(>F)
## race.location  3   0.63437 0.21146  1.1757 0.26074 0.2818
## Residuals     10   1.79864 0.17986         0.73926       
## Total         13   2.43301                 1.00000
```

```
source("Wd.R")
WdS.test(phy.dist, race.location, nrep = 99999)
```

```
## $p.value
## [1] 0.03777
## 
## $WdS.stat
## [1] 2.936692
## 
## $nrep
## [1] 99999
```

```
adonis(phy.dist~sample_data(physeq)$race, permutations=99999)
```

```
## 
## Call:
## adonis(formula = phy.dist ~ sample_data(physeq)$race, permutations = 99999) 
## 
## Permutation: free
## Number of permutations: 99999
## 
## Terms added sequentially (first to last)
## 
##                          Df SumsOfSqs MeanSqs F.Model      R2  Pr(>F)  
## sample_data(physeq)$race  1   0.34565 0.34565  1.9871 0.14207 0.06356 .
## Residuals                12   2.08736 0.17395         0.85793          
## Total                    13   2.43301                 1.00000          
## ---
## Signif. codes:  0 '***' 0.001 '**' 0.01 '*' 0.05 '.' 0.1 ' ' 1
```

```
WdS.test(phy.dist, sample_data(physeq)$race, nrep = 99999)
```

```
## $p.value
## [1] 0.04773
## 
## $WdS.stat
## [1] 2.157697
## 
## $nrep
## [1] 99999
```

```
adonis(phy.dist~sample_data(physeq)$location, permutations=99999)
```

```
## 
## Call:
## adonis(formula = phy.dist ~ sample_data(physeq)$location, permutations = 99999) 
## 
## Permutation: free
## Number of permutations: 99999
## 
## Terms added sequentially (first to last)
## 
##                              Df SumsOfSqs  MeanSqs F.Model      R2 Pr(>F)
## sample_data(physeq)$location  1   0.07524 0.075244 0.38296 0.03093 0.9073
## Residuals                    12   2.35777 0.196480         0.96907       
## Total                        13   2.43301                  1.00000
```

```
WdS.test(phy.dist, sample_data(physeq)$location, nrep = 99999)
```

```
## $p.value
## [1] 0.90838
## 
## $WdS.stat
## [1] 0.3829609
## 
## $nrep
## [1] 99999
```

Pairwise tests

```
with(sample_data(physeq), table(race, location))
```

```
##        location
## race    DIST PROX
##   black    2    3
##   white    5    4
```

All pairwise post-hoc tests

```
Tw2.posthoc.tests(phy.dist, race.location, nrep=999)
```

```
##      Level1       Level2       N1 N2 p.value tw2.stat  nrep
## [1,] "black.DIST" "white.DIST" 2  5  0.046   8.201647  999 
## [2,] "black.DIST" "black.PROX" 2  3  0.638   0.9802198 999 
## [3,] "black.DIST" "white.PROX" 2  4  0.061   2.922056  999 
## [4,] "white.DIST" "black.PROX" 5  3  0.63    0.832897  999 
## [5,] "white.DIST" "white.PROX" 5  4  0.456   0.984017  999 
## [6,] "black.PROX" "white.PROX" 3  4  0.949   0.5647803 999
```

1 vs All post-hoc tests

```
Tw2.posthoc.1vsAll.tests(phy.dist, race.location, nrep = 999)
```

```
##            N1 N2 p.value tw2.stat  nrep
## black.DIST 12 2  0.039   8.878768  999 
## white.DIST 9  5  0.075   1.926456  999 
## black.PROX 11 3  0.936   0.3560987 999 
## white.PROX 10 4  0.665   0.6978398 999
```
