## Supplementary material for "*W*_*d*_^*^-test: Robust Distance-Based Multivariate Analysis of Variance"

Analyze null simulation


### Analyze null simulation

###### *Alexander V. Alekseyenko*

#### *2/28/2017*

```
res = read.table('WStar_null.txt', header=T, row.names=1, sep="\t")
head(res)
```

```
##          n1 n2 n3 ef replicates Wstar Permanova    WelchF
## result.1  5  5  5  1          1 0.117     0.109 0.1135493
## result.2 10  5  5  1          1 0.878     0.900 0.8624573
## result.3 20  5  5  1          1 0.586     0.620 0.5918541
## result.4 40  5  5  1          1 0.373     0.635 0.3334131
## result.5  5 10  5  1          1 0.982     0.985 0.9729033
## result.6 10 10  5  1          1 0.851     0.845 0.8881348
```

```
Wstar.rr = with(res, tapply(Wstar, interaction(n1, n2, n3, ef), function(x) sum(x<0.05)))/1000
Permanova.rr = with(res, tapply(Permanova, interaction(n1, n2, n3, ef), function(x) sum(x<0.05)))/1000
WelchF.rr = with(res, tapply(WelchF, interaction(n1, n2, n3, ef), function(x) sum(x<0.05)))/1000

first= function(x) x[1]
n1.r = with(res, tapply(n1, interaction(n1, n2, n3, ef), first))
n2.r = with(res, tapply(n2, interaction(n1, n2, n3, ef), first))
n3.r = with(res, tapply(n3, interaction(n1, n2, n3, ef), first))
ef.r = with(res, tapply(ef, interaction(n1, n2, n3, ef), first))

plot.data = data.frame(n1.r, n2.r, n3.r, ef.r, Wstar.rr, Permanova.rr, WelchF.rr)

library(reshape2); packageVersion("reshape2")
```

```
## [1] '1.4.3'
```

```
ggdata = melt(plot.data, id.vars = c("n1.r", "n2.r", "n3.r", "ef.r"))
names(ggdata$value) = NULL
library(ggplot2); packageVersion("ggplot2")
```

```
## [1] '2.2.1'
```

```
tIe.plot = ggplot(ggdata, aes(y=value, x=n1.r, shape=factor(ef.r^2), color=variable)) + geom_point()+
  geom_line(aes(group=interaction(variable, ef.r^2)))+
  facet_grid(n2.r ~ n3.r, scales = "free_y")+theme_minimal() + 
  geom_hline(yintercept=0.05, color='red', lty='dashed')+
  scale_shape_discrete("Variance") + scale_color_discrete("Test") +
  ylab("Fraction of rejected null hypotheses") + xlab("n1") + 
  theme(legend.position = "top")+ guides(shape=guide_legend(title.position="top"),
                                        color=guide_legend(title.position="top"))
print(tIe.plot)
```

```
pdf('WdStartTypeI.pdf', width=6, height=6)
print(tIe.plot)
dev.off()
```

```
## quartz_off_screen 
##                 2
```

```
pval.plot = ggplot(res, aes(x = Wstar, y=WelchF)) + geom_point()+theme_minimal()
print(pval.plot)
```

```
pdf("WdStarVSWd.pdf",width=3, height=3)
print(pval.plot)
dev.off()
```

```
## quartz_off_screen 
##                 2
```

```
pval.perm=ggplot(res, aes(x = Permanova, y=WelchF)) + geom_point(aes(color=factor(ef^2)), size=0.3) +  theme_minimal() + scale_color_discrete("Variance") + theme(legend.position="bottom")
print(pval.perm)
```

```
pdf("WdStarVSPermanova.pdf",width=3, height=3)
print(pval.perm)
dev.off()
```

```
## quartz_off_screen 
##                 2
```
