## Supplementary material for "*W*_*d*_^*^-test: Robust Distance-Based Multivariate Analysis of Variance"

Load the functions


### Load the functions

```
source('Tw2.R')
ls()
```

```
## [1] "dist.cohen.d" "dist.sigma2"  "dist.ss2"     "is.dist"     
## [5] "Wstar"        "Wstar.test"   "WT"           "WT.mean"     
## [9] "WT.test"
```

### Univariate case test

In the univariate case, distance W\* is equivalent to the original Welch MANOVA implemented in R with `oneway.test`. The statistic for the test and distance W\* should be the same modulo numerical issues.

```
set.seed(2.15)

test.one = function(z){
  y=rnorm(100)
  dm = as.matrix(dist(y))
  f=factor(sample(3, 100, replace = T))
  Wstar(dm, f) / oneway.test(y~f)$statistic  
}

sapply(1:10, test.one)
```

```
## F F F F F F F F F F 
## 1 1 1 1 1 1 1 1 1 1
```

#### Compare theoretical results to W\* and PERMANOVA

##### Type I error

```
library(vegan); packageVersion("vegan")
```

```
## Loading required package: permute
```

```
## Loading required package: lattice
```

```
## This is vegan 2.4-2
```

```
## [1] '2.4.2'
```

```
simulate.null = function(ns, ef){
  y=rnorm(sum(ns), sd=rep(c(1,ef,ef^2), ns))
  f = factor(rep(c(1,2,3), ns))
  dm = as.matrix(dist(y))
  c(Wstar = Wstar.test(dm, f)$p.value, Permanova = adonis(dm~f)$aov.tab[1,6], WelchF = oneway.test(y~f)$p.value)
}
simulate.null(ns=c(5,5,5), ef=1)
```

```
##     Wstar Permanova    WelchF 
## 0.9900000 0.9890000 0.9901083
```

```
n.set = c(5,10,20,40)
ef = c(1, sqrt(0.2), sqrt(0.8))
params = expand.grid(n1=n.set, n2=n.set, n3=n.set,ef=ef, replicates=1:1000)
head(params)
```

```
##   n1 n2 n3 ef replicates
## 1  5  5  5  1          1
## 2 10  5  5  1          1
## 3 20  5  5  1          1
## 4 40  5  5  1          1
## 5  5 10  5  1          1
## 6 10 10  5  1          1
```

```
dim(params)
```

```
## [1] 192000      5
```

```
library(foreach); packageVersion("foreach")
```

```
## [1] '1.4.3'
```

```
library(doParallel); packageVersion("doParallel")
```

```
## Loading required package: iterators
```

```
## Loading required package: parallel
```

```
## [1] '1.0.10'
```

```
registerDoParallel(cores=8)

date()
```

```
## [1] "Mon Feb 27 14:30:59 2017"
```

```
res = foreach(i=1:(dim(params)[1]), .combine=rbind) %dopar% {
  c(params[i,], simulate.null(params[i,1:3], params[i,4]))
}
date()
```

```
## [1] "Mon Feb 27 23:54:08 2017"
```

```
#colnames(res) = c("Replicate", "n1", "n2", "fracSd", "effect", "adonis.p", "welchT.p", "welchTS.p")


write.table(res, "WStar_null.txt", sep="\t")
```
